# Transcriptomic and Biochemical analysis of *Procambarus clarkii* upon exposure to Pesticides: Population-Specific responses as a sign of pollutant resistance?

**DOI:** 10.1101/2024.10.08.617147

**Authors:** Martínez-Alarcón Diana, Celine Reisser, Montserrat Solé, Jehan-Hervé Lignot, Georgina Rivera-Ingraham

## Abstract

The effects that anthropogenic stressors may have on modulating species plasticity has been relatively unexplored; however, it represents a scientific frontier that may offer insights into their ability to colonize new habitats. To explore the advantage that inhabiting polluted environments may offer to invasive species, we selected the crayfish *Procambarus clarkii*, a species that can colonize and thrive in a wide range of aquatic environments, including heavily polluted ones. Here, we studied the molecular and physiological responses of crayfish when experimentally exposed to a pesticide mix of azoxystrobin and oxadiazon at sublethal concentrations. We compared these responses in three isolated crayfish populations in Southern France that are established in areas with different pollution levels: a) Camargue, seasonally affected by pesticide pollution; b) Bages-Sigean, impacted all year-round by domestic effluents and; c) Salagou, a more pristine site. Gene expression analyses revealed that the response to the pesticide mix was the strongest in the Camargue crayfish. In this population, a total of 88 differentially expressed genes (DEGs) were identified in hepatopancreas and 78 in gills between exposed and control laboratory groups. Among genes that were differentially expressed and successfully annotated, those involved in stress, DNA repair, immune response, and translation and transcription processes stand out. Interestingly, the hepatopancreas responded mainly with upregulation, but with downregulation in the gills. This suggests that compared to naïve individuals, when exposed to these biocides in their natural habitat crayfish respond with different mechanistic strategies that may confer them adaptability at the population level. Responses in terms of antioxidant and detoxification enzymes also corroborate differences to biocide inputs according to the origin of the crayfish.

## 1.1 Introduction

Exposure to harmful levels of chemical pollutants has led to the evolution and tolerance in populations of several marine species (Whitehead 2017; Reid et al. 2016; Hamilton et al. 2017; Oziolor et al. 2019). Although there is still a lack of evidence on the physiological basis for these adaptations, it is suspected that they may rely on processes involving absorption, distribution, and excretion of the chemical in question (Hamilton et al. 2017). While rapid evolution driven by pollutants has only been vaguely studied in aquatic invertebrates, research on killifish has provided insights into the key features enabling rapid evolutionary rescue in degraded environments. Studies analyzing killifish industrial pollutants exposure revealed impacts on nucleotide diversity and several molecular structures (Reid et al. 2016; Whitehead et al. 2017; Hamilton et al. 2017), and multiple metabolic pathways (Reid et al. 2016; Oziolor et al. 2019). Potential adaptive mechanisms include enhanced antioxidant responses and increased capacity for DNA and tissue repair (Hamilton et al. 2017). These findings suggest that the evolutionary influence of anthropogenic stressors, as selective agents, is a widespread phenomenon (Whitehead et al. 2017). Nonetheless, there is very limited information on how these stressors may affect the tolerance, adaptation, and rapid evolution of aquatic invertebrates. Here, we address pesticides adaptation of aquatic invertebrates using the crayfish *Procambarus clarkii* as a study model. This invasive species is well known for its ability to colonize a wide range of aquatic environments with different levels of water quality. It can disperse widely, tolerate environmental extremes, has generalist and opportunistic feeding habits (Gherardi and Barbaresi, 2007), and is more resistant to diseases than most of its native counterparts (Collas et al. 2007). It is native to the North of Mexico and the United States, but its highly adaptive nature has driven it to be well-established throughout Europe, Asia, Africa, North America and South America. It was first introduced in Western France in 1974, and by the mid-1990s it had established populations in 36 of the 96 metropolitan France counties (departments), particularly around the coastal Mediterranean areas (Meineri et al. 2013). Previous studies with this species showed that responses to specific pesticides differed between individuals from populations coming from more pristine or polluted environments. Some of these differences included respiration rates, hydro-osmotic balance, and the activity of digestive proteases and lipases (Raffalli et al. 2024). In this study, we analyzed and compared the gene expression and enzymatic responses in gills and hepatopancreas (midgut) of the same populations following a 96-hour laboratory exposure to a pesticide cocktail containing azoxystrobin and oxadiazon at sublethal concentrations.

In the last decades, most studies in ecotoxicology have focused on the analysis of well-known biochemical parameters to measure the impact of toxicants on organisms. However, the impact may extend beyond these stress mechanisms, and by limiting the focus of the research, we could be missing the bigger picture. For this reason, in this study, we analyzed the entire set of genes that are transcribed in response to toxicant exposure. We investigated whether gene expression in the hepatopancreas and gills differ between these three populations and which were most impacted pathways upon laboratory exposure to the pesticide mix. Furthermore, to complement gene expression data, we targeted antioxidant and detoxification enzymatic responses by monitoring the activity of physiological markers that are good stress indicators in marine and aquatic invertebrates. We focused our analyses on gills and hepatopancreas due to their role in crustacean toxicology: gills are the primary organ of respiration and osmoregulation, and the first barrier of exposure to water-borne chemicals (Burnett et al. 1985) whereas the hepatopancreas that is part of the digestive system, another major entry route for toxicants, plays a major role in detoxification processes (White and Rainbow 1986; Liu X et al. 2021). For this purpose, crayfish were collected from three populations in the South of France: i) the brackish Camargue wetland system, seasonally affected by pesticide pollution; ii) the brackish Bages-Sigean lagoons, regularly impacted by pesticides together with domestic effluents and; iii) the Salagou lake, a more pristine freshwater site. We hypothesized that crayfish subjected in a life-long manner to varying pulses of environmental chemicals would exhibit different responses to additional pollutant stressors. Consequently, we anticipated that the Camargue and Bages-Sigean populations would show a distinct gene expression fingerprint compared to the Salagou one, which likely had not previously faced exposure to these pollutants. Furthermore, we also hypothesized that the antioxidant and detoxification responses would be lower in populations that formerly and repeatedly encounter pesticides in their natural environment.

## 2. Methods

### 2.1 Animal sampling and maintenance

Individuals from 3 populations in the South of France were collected using land nets from: i) the Fumemorte canal in the Camargue area (43°30’52.6”N, 4°40’02.1”E), with an environmental salinity ranging 24-33 g/L and degraded environmental quality from March to September, due to the intense agricultural activity primarily consisting of rice fields in the area, and hence, pesticide pollution; ii) Bages-Sigean wetlands (43°07’30.7”N, 3°01’20.9”E) (from Bages-Sigean region), an area with a salinity of 26-40 g/L and presenting degraded water quality throughout the year, associated to urban and domestic discharges and; iii) Salagou lake (43°39’45.0”N, 3°22’20.5”E), a freshwater body <1 g/L presenting an overall good environmental quality (Fig. 1).

**Fig. 1.**
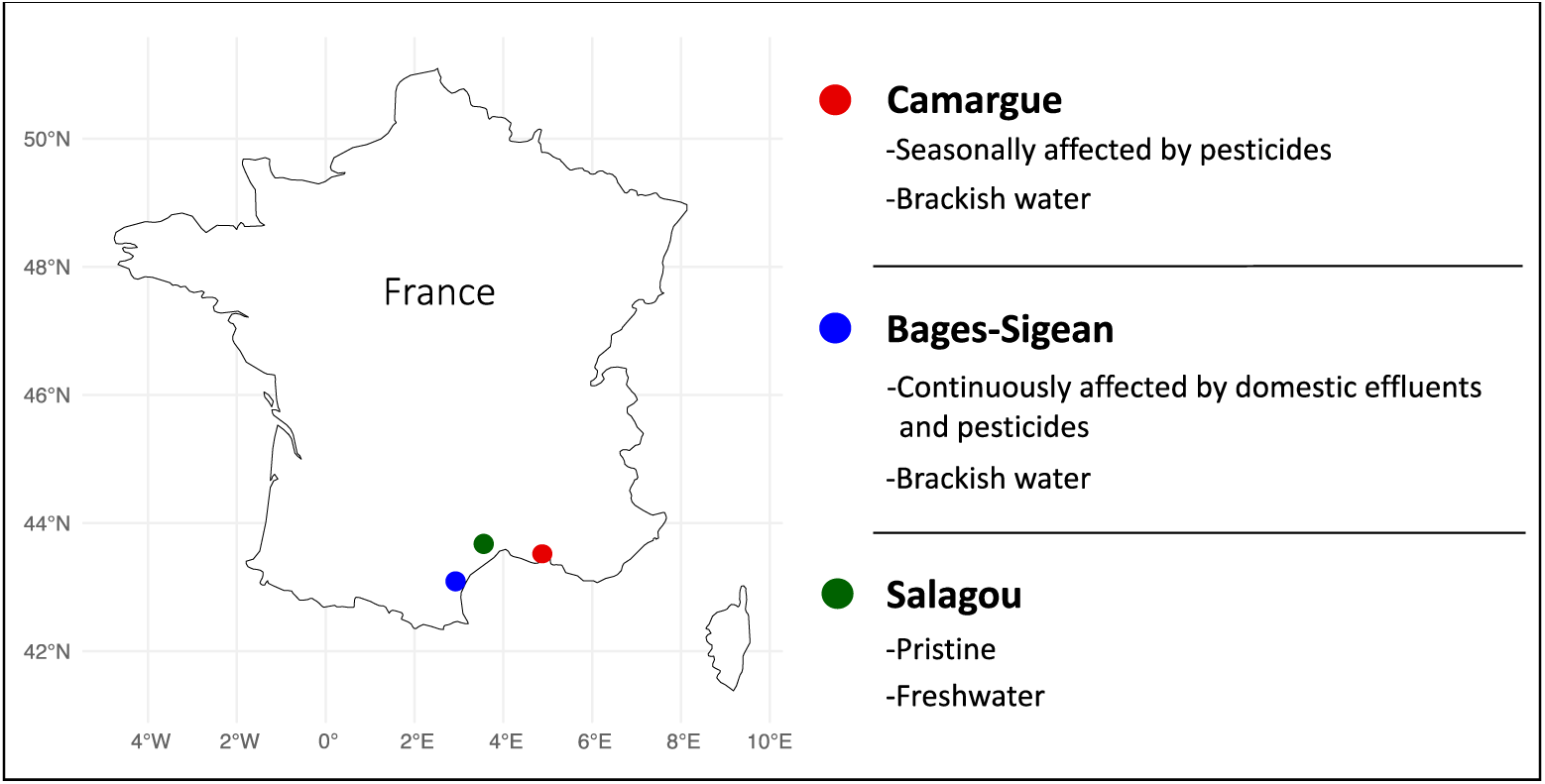
Sampling areas of crayfish, *Procambarus clarkii* analyzed in this study. The color of the circles represents the populations: Red for Camargue, blue for Bages-Sigean and green for Salagou.

### 2.2 Experimental design

After field collection, animals were transported to the laboratory where they were acclimated to freshwater for 4 months in large aquaria. A total of 90 individuals (1/1 sex ratio) were used for this study: 15 individuals (5 per population) were distributed among six tanks in 20-L aquaria. Within the aquaria, each individual was placed in an immersed glass jar, closed by a glass wire net to individually identify each animal and avoid the negative impact of social conflicts among them. Animals were allowed to acclimate to the new housing conditions (i.e. the glass jars) for 1 week before the start of the experiments. After this time, half of the animals remained undisturbed for control purposes while the other half were exposed to a mixture of 2 pollutants: azoxystrobin (95 µg/L, Sigma-Aldrich 31697) and oxadiazon (30 µg/L, Sigma-Aldrich 33382) for 96h. These concentrations are 100 times higher than the maximum acceptable concentrations according to the environmental quality standards for these two pesticides (EFSA 2010). Because a preliminary test determined that the concentration of the pesticides decreased around 10% in 24h, a daily renew of 80% of the water along with the corresponding doses of pollutants was done to ensure a constant chemical concentration throughout the 4-day experimental time. The chemicals were diluted in methanol for their delivery in the corresponding aquaria, while controls received the same amount of methanol. The total concentration of methanol in each aquarium was 0.018 ml/L, which is below the lowest concentration for causing effect due to chronic exposure to methanol (Kaviraj et al. 2004). During crayfish acclimation to laboratory conditions and during the biocide exposures, water temperature was maintained at 20°C (±0.4°C) with a photoperiod of 12:12-hour. All animals were maintained in recirculated and dechlorinated tap water.

### 2.3 Sample collection

After the 96h-exposure, animals were removed from the aquaria and euthanized following RSPCA policies: here, crayfish were immediately anaesthetized through air-chilling method, i.e. placing the animals in a −20°C chamber for 15 minutes, to make them insensitive to stimuli. After this time, animals were euthanized by removal of the frontal part of the cephalothorax and the tissues dissected. Hepatopancreas and gills samples were flash-frozen in liquid nitrogen and stored −80°C for enzyme analyses. They were aslo stored in microtubes containing RNAlater® Stabilization Solution (Ambion, Inc., Texas, USA) and posteriorly stored at −80°C for gene expression analyses. All experiments were conducted in accordance with the valid international, European and National laws, applying the principles of replacement, reduction and refinement.

### 2.4 RNA extraction

About 30mg of hepatopancreas and gill tissue previously stored at −80°C in RNAlater® were used for RNA extraction. Cell lysis was performed in 0.6 ml of lysis buffer RLT provided in the Qiagen Kit (Qiagen, Hilden, Germany) with beta-mercaptoethanol and using ceramic beads (Precellys® Keramik-kit, PeqLab, Erlangen, Germany), with 16 seconds shaking at 6500 rpm. Subsequently, samples were centrifuged at 16,000 g for 10 minutes and then transferred into a 2 ml microtube. Total RNA was isolated using RNAeasy Mini spin columns (Qiagen, Germany) following the manufacturer’s instructions. RNA quantity was analyzed using a NanoDrop One spectrophotometer (Thermo Fisher Scientific) and quality was determined by microfluidic electrophoresis in a Bioanalyzer (Agilent Technologies, USA).

### 2.5 RNAseq normalized cDNA libraries and Ilumina sequencing

The construction of cDNA libraries from 64 individuals was done by Macrogen Europe (Amsterdam, Netherlands) following the TruSeq stranded mRNA sample protocol. To verify the size of PCR enriched fragments, the template size distribution was checked by running on an Agilent Technologies 2100 Bioanalyzer using a DNA 1000 chip. Additionally, the prepared libraries were quantified using qPCR according to the Illumina qPCR Quantification Protocol Guide. Finally, to calculate the library sample concentration we also used Roche’s Rapid library standard Quantification solution and calculator. Only libraries with a concentration over 10 nM were used. Subsequently, the libraries were sequenced on an Illumina NovaSeq 6000 sequencer by Macrogen Europe with a throughput of 80M reads per sample (100bp paired-end).

### 2.6 Differential gene expression analysis and functional annotation

Trimmomatic software in paired-end mode was used for quality filtering of the raw RNA reads and adapter filtering. Reads with a quality below 28, and a length less than 40bp were discarded. After the filtering process, clean reads were mapped onto *P. clarkii* genome assembly (Xu et al. 2021), using bwa-mem2 (Vasimuddin et al. 2019) with default parameters. Alignments were then filtered to discard unmapped reads, and we used featureCounts (Liao et al. 2014) to obtain, a table of read count per transcript.

Functional annotation of the differentially expressed transcripts was performed using the fasta protein file of the transcripts from the genome, and the BeeDeeM pipeline (https://github.com/pgdurand/BeeDeeM) on Uniprot and Swissprot databases.

### 2.7 Biochemical analyses

A portion of the hepatopancreas and gills were homogenized in 100 mM phosphate buffer in a 1:5 (w:v) ratio using the Precellys keramik-kit (MP, Germany), performing 2 cycles of 15 seconds shaking and a 20 seconds pause in between. The buffer used for hepatopancreas (100 mM phosphate buffer) also contained 150 mM KCl and 1mM EDTA. The homogenates were centrifuged at 10,000g for 20 minutes at 4°C and the resulting supernatants were stored at −80°C for further biochemical determinations. Protein quantification was performed after Bradford (1976) with bovine serum albumin as standard (A9418, Sigma Aldrich).

Antioxidant capacity was estimated through the catalase (CAT) and glutathione reductase (GR) enzyme activities. Detoxification and biotransformation were assessed as glutathione-*S*-transferase (GST) and carboxylesterase (CE) enzyme activities. CAT activity was measured in hepatopancreas at 240 nm following the method described by Aebi (1984). GR activity in hepatopancreas supernatants was measured at 340 nm during 3 min adapted from the method described by Carlberg and Mannervik (1985). GST activity was measured in both hepatopancreas and gills homogenates at 340 nm for 3 min following the method described by Habig et al. (1974). Carboxylesterase (CE) activity in hepatopancreas and gills was measured using p-nitrophenyl acetate p-NPA (N8130, Sigma Aldrich) and p-nitrophenyl butyrate p-NPB (N9876, Sigma) as substrate at 405 nm for 3 min following the protocol described by Hosokawa and Satoh (2001). Furthermore, neurotoxicity assessment was measured as acetylcholinesterase (AChE) and the potential to interfere with molting activity by means of N-acetyl-β-D-glucosaminidase (NAGase) activities. The inhibition of AChE activity is one of the most frequently adopted biomarkers for neurotoxicity by pesticides but also other environmental chemicals (see review by Fu et al. 2018) and exploited for pest control monitoring purposes (Casida and Durkin, 2013; Lignot et al. 1998). Environmental chemicals such heavy metals (e.g. Zhang et al. 2008; Han and Wang, 2009; Rivera-Ingraham et al. 2021), hydrocarbons (Zhang et al. 2008), pharmaceuticals (e.g. Rhee et al. 2013) or even herbicides like the ones here tested (e.g. Kovačević et al. 2023) inhibit AChE. While NAGase has been suggested as a good biomarker of molting toxicity as it is affected by a wide variety of environmental pollutants (e.g. Lin et al., 2005; Zhang et al., 2010; Mesquita et al., 2015).

AChE was measured in hepatopancreas and gills by using acetylthiocholine as substrate and the kinetics of the metabolite formed with DTNB was read at 412nm for 5 min following the protocol described by Ellman et al., (1961). NAGase activity was determined using 4-nitrophenyl N-acetyl-β-D-glucosaminide as substrate and spectrophotometrically recording the formation of 4-nitrophenol at 410 nm also for 5 min (Rollin et al. 2023).

All enzymatic activities were expressed per mg of protein content, as measured using the Bradford (1976) method and bovine serum albumin (BSA) (0.05-0.5 mg/mL) as standard. All activities were measured with an Infinite200 TECAN spectrophotometer (Tecan, Männendorf, Switzerland) at 25 °C using the Magellan kinetic mode v6.0.

### 2.8 Statistical analyses

For statistical analyses of enzymatic activities, RStudio version 2023.03.0 software was used. Shapiro’s test was used to verify normal distribution and homogeneity of variances of biochemical data. One-way ANOVAs were used to compare populations within each condition. Differences among groups were subsequently identified by pairwise comparisons using Tukey’s post-hoc test. To test difference among treatments and controls from every population the non-parametric Wilcox test was applied. The level for statistical significance was set at P < 0.05. Differential gene expression was assessed using the Bioconductor R package DESeq2 (Love et al., 2014), with an alpha of 0.001.

## 3. Results

### 3.1 Differential gene expression

A multidimensional Principal Components Analysis (PCA) based on the gene expression analysis of the three crayfish populations clearly identified hepatopancreas and gill tissues in specific clusters (Fig 2A), with PC1 explaining 72.85% of the variance. Population’s dispersion can be observed in PC2, PC3, and PC4 which altogether explained 7.2% of the variance (Fig 2A, B).

**Figure 2.**
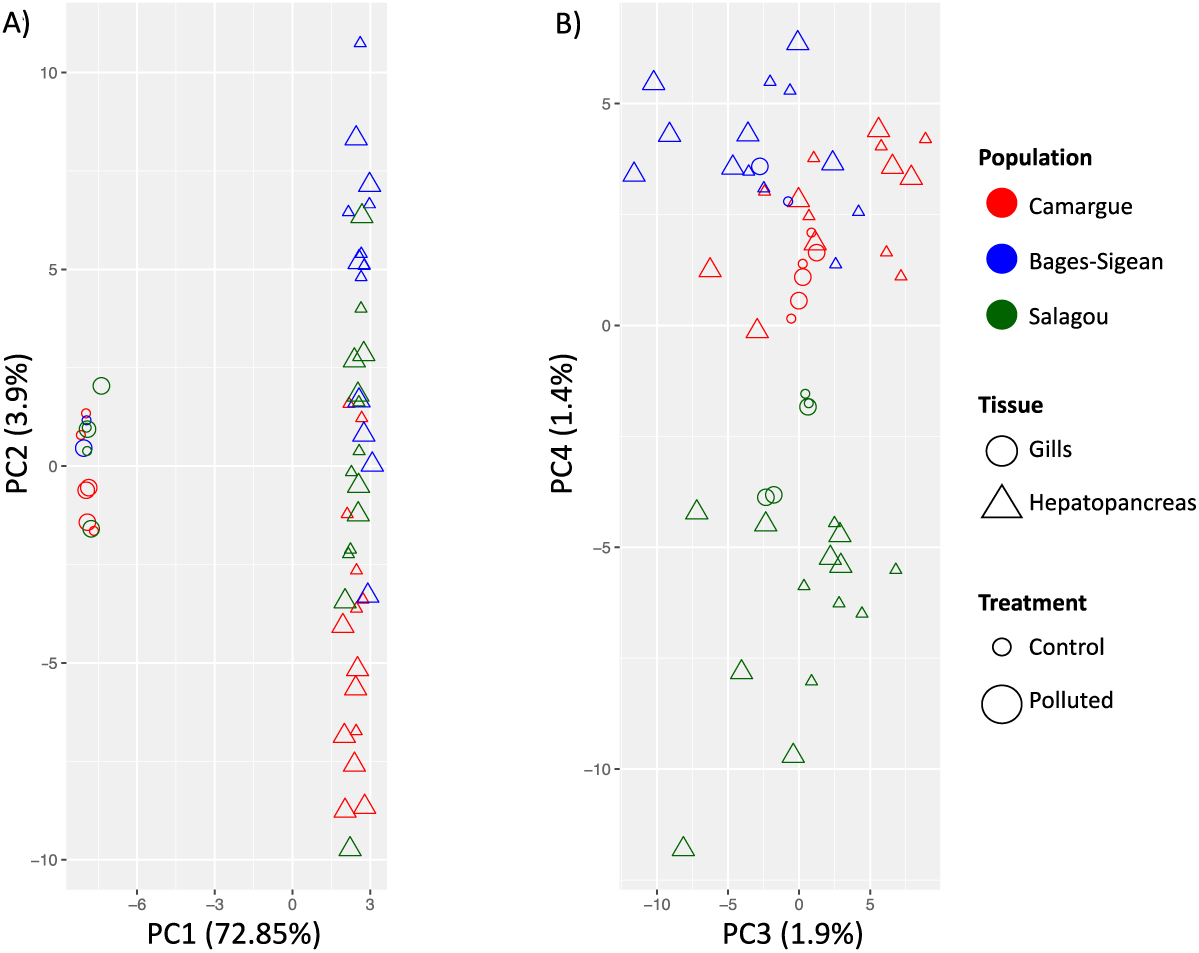
PCA of differentially expressed genes between tissues and populations. Circles and triangles represent gill and hepatopancreas results, respectively, and their size differs according to treatment (smaller for control animals and larger for pollutant-exposed animals). Different colors represent the three different populations considered: red for Camargue, blue for Bages-Segean, and green for Salagou.

In the hepatopancreas, a heatmap analysis of the differentially expressed genes (DEG) corroborated that when comparing all three populations simultaneously, the difference between lab-exposed and unexposed crayfish individuals is not evident (Fig. 3A). However, when analyzing each population separately, we observed a clear difference upon exposure to the pesticide mixture, with the most distinct pattern in Camargue crayfish (Fig. 3B, C, D). In this tissue, DEG under the pesticide mixture also differed among populations: Camargue showed 88 DEG (52 up-regulated and 36 down-regulated genes), far exceeding the changes experienced by Bages-Sigean crayfish (5 up-regulated and 4 down-regulated) and Salagou ones (6 up-regulated and 1 down-regulated) (Fig. 3C, D). However, not all DEGs were functionally annotated: only 55 genes out of 88 in Camargue, 6 out of the 7 in Salagou and 7 out of the 9 in Bages-Sigean.

**Figure 3.**
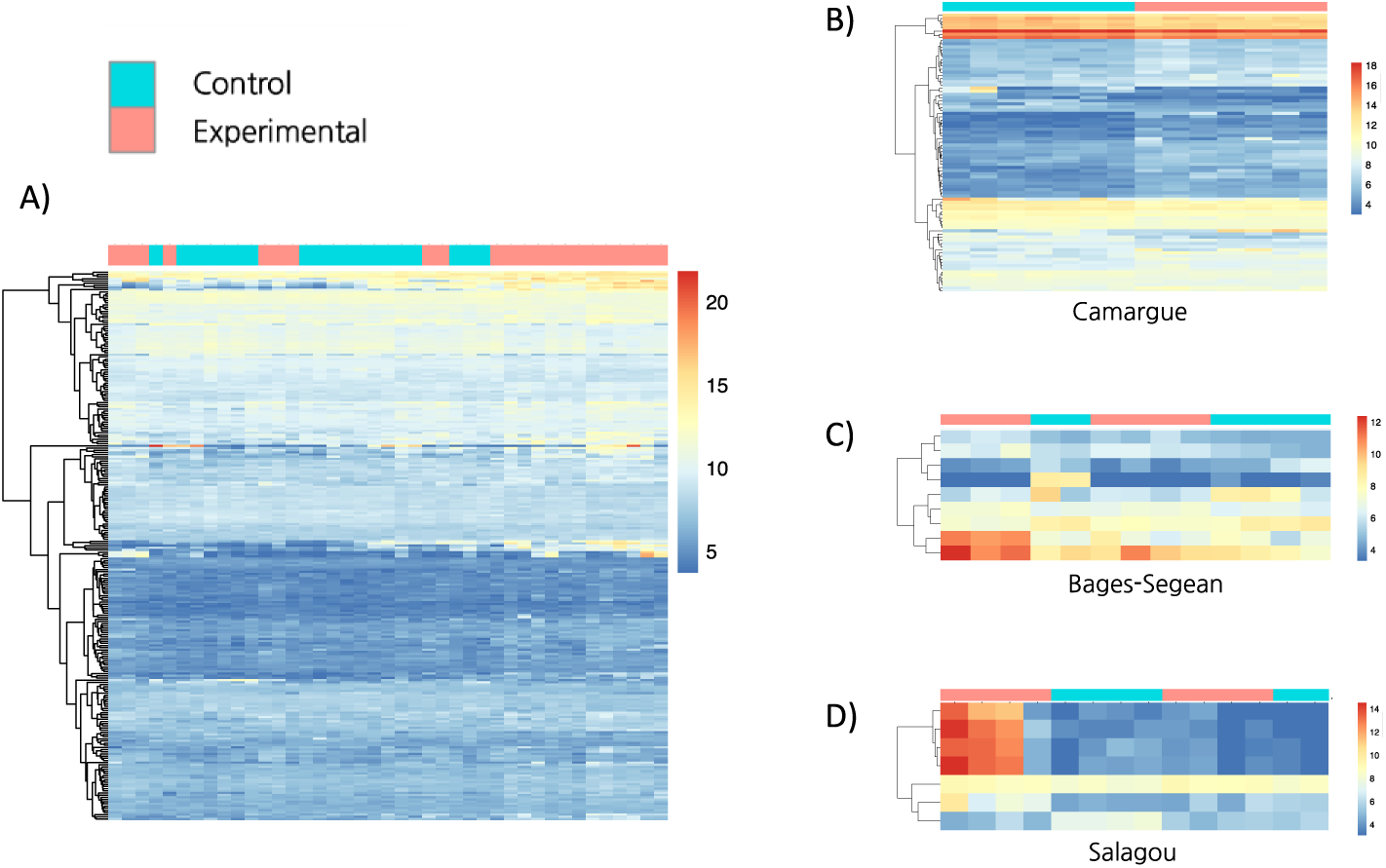
Heatmaps showing differentially expressed genes (DEGs) in the hepatopancreas of crayfish after 96-hour pesticide mixture exposure in animals from the three studied populations: A) General overview; B) Camargue; C) Bages-Sigean; D) Salagou.

For the particular case of Camargue crayfish, among the genes that were differentially expressed in the hepatopancreas and successfully annotated, those involved in stress response (6 genes up- and 4 down-regulated), DNA repair (5 up and 2 down), immune response (8 up and 2 down), and translation and transcription processes (4 up and 3 down) stand out (Table 1). In contrast to Camargue crayfish, the Salagou specimens upregulated only two genes related to stress and immune response in the hepatopancreas (Table 1).

**Table 1:** List of regulated genes involved in stress response, DNA repair, immune response and translation and transcription according to tissue and crayfish population. Positive values indicate upregulation, and negative values indicate downregulation.

| Hepatopancreas |  |  |  |
| --- | --- | --- | --- |
| Transcript | log2FoldChange | Type of response |  |
| Insulin-like growth factor-binding protein 7 | +1.259 | Stress response & DNA repair | GO:0034599: Cellular response to oxidative stress |
|  |  |  | GO:0006977: DNA damage response |
| Heat shock 70 kDa protein cognate 4-like | -1.036 | Stress response | GO:0009408: Response to heat |
|  |  |  | GO:0034620: Cellular response to unfolded protein |
|  |  |  | GO:0034976: Response to endoplasmic reticulum stress |
| Hemocyte protein-glutamine gamma-glutamyltransferase-like | +1.507 | Stress response | GO:0009611: Response to wounding |
|  |  |  | GO:0050832: Defense response to fungus |
| LOW QUALITY PROTEIN: nitric oxide synthase-like protein | +2.481 | Stress response | GO:0034614: Cellular response to reactive oxygen species |
|  |  |  | GO:0006979: Response to oxidative stress |
| Thioredoxin-like protein 1 | -0.654 | Stress response & DNA repair | GO:0034599: Cellular response to oxidative stress |
|  |  |  | GO:0006281: DNA repair |
|  |  |  | GO:0006979: Response to oxidative stress |
| DNA damage-inducible transcript 4-like protein | +3.56 | Stress response & DNA repair | GO:0006974: DNA damage response |
|  |  |  | GO:0006281: DNA repair |
|  |  |  | GO:0033554: Cellular response to stress |
| Protein spaetzle 4-like | -2.313 | Stress response | GO:0006952: Defense response |
|  |  |  | GO:0009611: Response to wounding |
| Heat shock cognate 70 kDa protein | -1.6 | Stress response | GO:0009408: Response to heat |
|  |  |  | GO:0034620: Cellular response to unfolded protein |
|  |  |  | GO:0034976: Response to endoplasmic reticulum stress |
| Vascular endothelial growth factor receptor 1-like | +1.744 | Stress response | GO:0008285: Negative regulation of cell proliferation |
|  |  |  | GO:0007169: Transmembrane receptor protein tyrosine kinase signaling pathway |
| Chorion peroxidase-like | +1.746 | Stress response | GO:0006979: Response to oxidative stress |
|  |  |  | GO:0033554: Cellular response to stress |
| Histone H1-delta-like | +1.259 | DNA repair | GO:0006974: DNA damage response |
|  |  |  | GO:0006303: Double-strand break repair via non-homologous end joining |
| SUMO-activating enzyme subunit 1-like | -0.505 | DNA repair | GO:0006302: Double-strand break repair |
|  |  |  | GO:0006301: Postreplication repair |
| DNA-directed RNA polymerase II subunit RPB1-like | +1.398 | DNA repair | GO:0006974: DNA damage response |
|  |  |  | GO:0006281: DNA repair |
| E3 ubiquitin-protein ligase TRIM9-like | +0.521 | DNA repair | GO:0043161: Proteasome-mediated ubiquitin-dependent protein catabolic process |
| Folate receptor beta-like | -1.889 | Immune response | GO:0050776: Regulation of immune response |
|  |  |  | GO:0009408: Response to heat |
| Astakine-like | +1.577 | Immune response | GO:0006955: Immune response |
|  |  |  | GO:0045087: Innate immune response |
| Beta-1,3-glucan-binding protein-like | +2.115 | Immune response | GO:0008233: Peptidase activity |
|  |  |  | GO:0050830: Defense response to Gram-positive bacterium |
|  |  |  | GO:0006952: Defense response |
| Leukocyte elastase inhibitor-like | +2.57 | Immune response | GO:0004866: Endopeptidase inhibitor activity |
|  |  |  | GO:0009611: Response to wounding |
|  |  |  | GO:0050727: Regulation of inflammatory response |
| Pulmonary surfactant-associated protein D-like | +1.321 | Immune response | GO:0006955: Immune response |
|  |  |  | GO:0008009: Chemokine activity |
|  |  |  | GO:0030203: Glycosaminoglycan metabolic process |
| Integrin alpha-8-like | +1.300 | Immune response | GO:0007229: Integrin-mediated signaling pathway |
|  |  |  | GO:0050900: Leukocyte migration |
| CLIP domain-containing serine protease 2-like | +1.741 | Immune response | GO:0008236: Serine-type peptidase activity |
|  |  |  | GO:0006955: Immune response |
|  |  |  | GO:0006952: Defense response |
| Angiotensin-converting enzyme-like | +7.521 | Immune response | GO:0002003: Angiotensin maturation |
|  |  |  | GO:0006955: Immune response |
| Protein spaetzle 4-like | -2.313 | Immune response | GO:0008063: Toll signaling pathway |
|  |  |  | GO:0006955: Immune response |
|  |  |  | GO:0006952: Defense response |
| Vascular endothelial growth factor receptor 1-like | +1.744 | Immune response | GO:0001525: Angiogenesis |
|  |  |  | GO:0050900: Leukocyte migration |
| Translation elongation factor 2-like | -0.609 | Translation and transcription | GO:0003746: Translation elongation factor activity |
| Eukaryotic translation initiation factor 3 subunit B-like | -0.741 | Translation and transcription | GO:0003743: Translation initiation factor activity |
| DNA-directed RNA polymerase II subunit RPB1-like | +1.398 | Translation and transcription | GO:0006351: DNA-templated transcription |
| Transcription initiation factor TFIID subunit 1-like | +2.806 | Translation and transcription | GO:0006352: DNA-templated transcription initiation |
| Mediator of RNA polymerase II transcription subunit 19-like | -0.681 | Translation and transcription | GO:0006351: DNA-templated transcription |
| Elongator complex protein 6-like isoform X1 | +2.100 | Translation and transcription | GO:0003723: RNA binding |
| mRNA decay activator protein ZFP36L1-like isoform X1 | +0.693 | Translation and transcription | GO:0006402: mRNA catabolic process |
| **Gills** |  |  |  |
| Endochitinase-like | -8.02 | Stress response & Immune response | GO:0009409: Response to cold |
|  |  |  | GO:0006032: Chitin catabolic process |
|  |  |  | GO:0008063: Toll signaling pathway |
|  |  |  | GO:0050832: Defense response to fungus |
| Proline-rich extensin-like protein EPR1 | -1.703 | Stress response | GO:0006950: Response to stress |
|  |  |  | GO:0009651: Response to salt stress |
| E3 ubiquitin-protein ligase RNF12-B-like | -4.486 | Stress response & DNA repair | GO:0006950: Response to stress |
|  |  |  | GO:0006281: DNA repair |
|  |  |  | GO:0006974: DNA damage response |
|  |  |  | GO:0016567: Protein ubiquitination |
| Chitin deacetylase 1-like | -3.442 | Stress response & Immune response | GO:0009409: Response to cold |
|  |  |  | GO:0006952: defense response |
|  |  |  | GO:0008063: Toll signaling pathway |
|  |  |  | GO:0010200: Response to chitin |
| Transcription initiation factor TFIID | -6.405 | DNA repair | GO:0003684: Damaged DNA binding |
|  |  |  | GO:0006352: DNA-templated transcription initiation |
|  |  |  | GO:0005669: Transcription factor TFIID complex |
| Cyclin-dependent kinase inhibitor | -1.414 | DNA repair | GO:0003684: Damaged DNA binding |
|  |  |  | GO:0006281: DNA repair |
|  |  |  | GO:0051726: Cellular cycle regulation |
|  |  |  | GO:0006974: DNA damage response |
| Lysosomal aspartic protease-like | -6.241 | Immune response | GO:0004190: Aspartic-type endopeptidase activity |
|  |  |  | GO:0006915: Apoptotic process |
|  |  |  | GO:0006952: Defense response |
| Protein spaetzle 4-like | -1.059 | Immune response | GO:0008063: Toll signaling pathway |
|  |  |  | GO:0006955: Immune response |
|  |  |  | GO:0006952: Defense response |
| Anti-lipopolysaccharide factor-like | -1.745 | Immune response | GO:0006952: Defense response |
| Transcription initiation factor TFIID subunit 13-like isoform X3 | -6.405 | Translation and transcription | GO:0006352: DNA-templated transcription initiation |
|  |  |  | GO:0006367: Transcription initiation at RNA polymerase II promoter |
| Zinc finger protein 271-like | +1.054 | Translation and transcription | GO:0006351: DNA-templated transcription |
|  |  |  | GO:0003677: DNA binding |
| Cyclin-dependent kinase inhibitor 1-like | -1.414 | Translation and transcription | GO:0003676: Nucleic acid binding |
|  |  |  | GO:0006355: Regulation of DNA-templated transcription |
| Heterogeneous nuclear ribonucleoprotein A3 homolog 1-like | -2.364 | Translation and transcription | GO:0003676: Nucleic acid binding |
|  |  |  | GO:0000398: mRNA splicing, via spliceosome |
| Translation initiation factor IF-2-like | -3.940 | Translation and transcription | GO:0006413: Translational initiation |
| Ribosome-binding protein 1-like | -1.256 | Translation and transcription | GO:0006412: Translation |
| Elongation factor 1-alpha-like | +4.369 | Translation and transcription | GO:0006414: Translational elongation |

Salagou Population
| **Hepatopancreas** |  |  |  |
| --- | --- | --- | --- |
| Transcript | log2FoldChange | Type of response according to enrichment analysis |  |
| Delta-1-pyrroline-5-carboxylate synthase-like | +3.481 | Stress response | GO:0003842: 1-pyrroline-5-carboxylate dehydrogenase activity. |
| Pseudohemocyanin-2-like (4 isoforms) | +10.50 | Immune response | GO:0006952: Defense response |
|  | +9.29 |  | GO:0019826: Oxygen transport |
|  | +11.45 |  |  |
|  | +10.342 |  |  |
| **Gills** |  |  |  |
| Calmodulin-like: | +2.10 | Stress response | GO:0005509: Calcium ion binding |
|  |  |  | GO:0005516: Calmodulin binding |
| Digestive cysteine proteinases | +26.14 | Stress response | GO:0006508: Proteolysis |
|  |  |  | GO:0006952: Defense response |
| WAP four-disulfide core domain protein 2-like | +2.107 | Immune response | GO:0006952: Defense response |
| Techylectin-5A-like | -1.898 | Immune response | GO:0006955: Immune response |
|  |  |  | GO:0050832: Defense response to fungus |
| Mucin-17-like | -2.070 | Immune response | GO:0006952: Defense response |
|  |  |  | GO:0007165: Signal transduction |
| Probable chitinase 10 | +8.229 | Immune response | GO:0006032: Chitin catabolic process |
|  |  |  | GO:0006952: Defense response |
| Chitinase-3-like protein 1 | +6.457 | Immune response, | GO:0006032: Chitin catabolic process |
|  |  |  | GO:0006952: Defense response |
| Hematopoietic prostaglandin D synthase-like (2 ISOFORMS) | +4.503<br>+2.383 | Immune response | GO:0006693: Prostaglandin metabolic process |
|  |  |  | GO:0006952: Defense response |
| Protein Skeletor, isoforms B/C-like isoform X2 | +1.450 | Translation and transcription | GO:0006351: DNA-templated transcription |
|  |  |  | GO:0006355: Regulation of DNA-templated transcription |

Bages-Sigean Population
| Hepatopancreas |  |  |  |
| --- | --- | --- | --- |
| Transcript | log2FoldChange | Type of response according to enrichment analysis |  |
| Daf-12-interacting protein 1-like | +1.656 | Stress response | GO:0009267: Cellular response to starvation |
|  |  |  | GO:0042594: Response to starvation |
|  |  |  | GO:0033554: Cellular response to stress |
|  |  |  | GO:0009628: Response to abiotic stimulus |
| Arylalkylamine N-acetyltransferase-like 2 | -3.022 | Stress response | GO:0006970: Response to osmotic stress |
|  |  |  | GO:0007585: Respiratory gaseous exchange by respiratory system |
|  |  |  | GO:0007623: Circadian rhythm |
| Facilitated trehalose transporter Tret1-2 homolog | +1.563 | Stress response | GO:0006970: Response to osmotic stress |
|  |  |  | GO:0009651: Response to salt stress |
|  |  |  | GO:0042538: Hyperosmotic salinity response |
| Origin recognition complex subunit 3-like | -1.091 | DNA repair | GO:0006270: DNA replication initiation |
|  |  |  | GO:0003688: DNA replication origin binding |
|  |  |  | GO:0031297: Replication fork processing |
| Low density lipoprotein receptor adapter protein 1-A-like | +0.793 | Immune response | GO:0006955: Immune response |
| Origin recognition complex subunit 3-like | -1.091 | Translation and transcription | GO:0003677: DNA binding |
|  |  |  | GO:0006355: Regulation of DNA-templated transcription |
|  |  |  | GO:0006270: DNA replication initiation |

Contrarily to the results obtained for the hepatopancreas, differential gene expression in gills can more easily separate the experimentally exposed and control individuals (Fig. 4A). As observed in hepatopancreas, the gills of Camargue crayfish exhibited a higher number of DEG under pesticide mixture exposure in comparison with the unexposed controls (Fig. 4A).

**Fig. 4.**
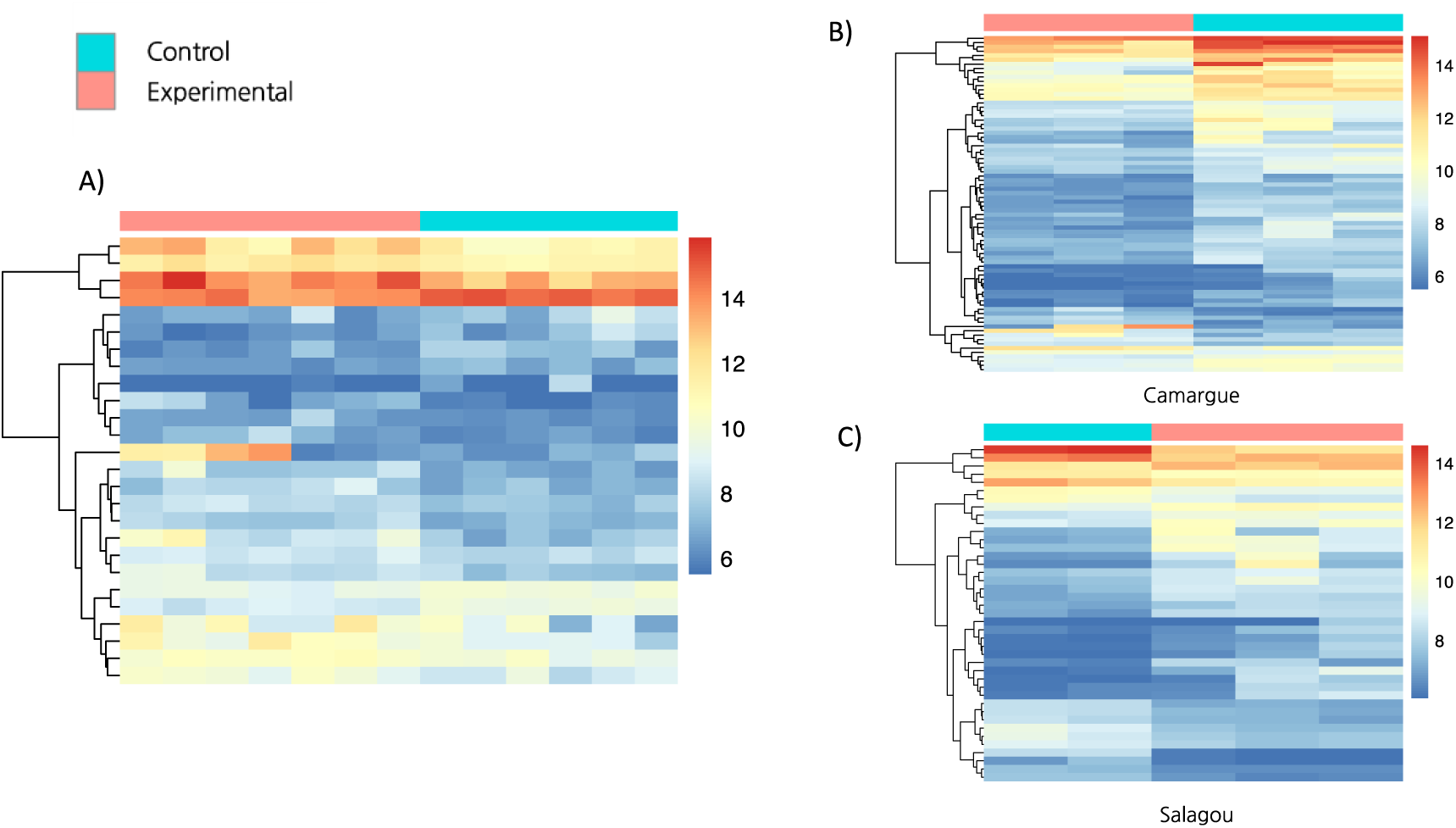
Heatmaps showing differentially expressed genes (DEGs) in gills of crayfish after 96-hour pesticide mixture exposure in animals from the three studied populations: A) General overview, and from two of the studied populations B) Camargue; C) Salagou.

Out of the 78 DEGs identified in gills of Camargue crayfish, 55 were annotated. Among the 78 DEGs, 67 genes were down-regulated. Genes involved in stress response, DNA repair, immune response, and translation and transcription were mostly down-regulated, except for a zinc finger protein and elongation factor, which were upregulated and are involved in translation and transcription (Table 1). For crayfish from Bages-Sigean no DEG analysis was possible, since only 2 samples met the library quality requirements for sequencing.

### 3.2 Biochemical assessments: antioxidant and detoxification activities

Enzymatic results are summarized in Table 2. Among all the enzymatic determinations carried out, statistical differences between controls and pesticide-exposed animals were detected in hepatopancreas (Fig. 5D, 5F, 5G) but not in gills (Fig. 6). Neither CE activities measured with both substrates (pNPA or pNPB) nor AChE were affected by the 96h treatment in either population; although, on average, hydrolysis rates in hepatopancreas were 80% higher than in gills (Fig. 5C and 6C), as it corresponds to the main metabolic organ. A higher metabolic responsiveness in hepatopancreas than in gills coincides with the enhanced transcriptomic changes also observed in the hepatic tissue.

**Figure 5.**
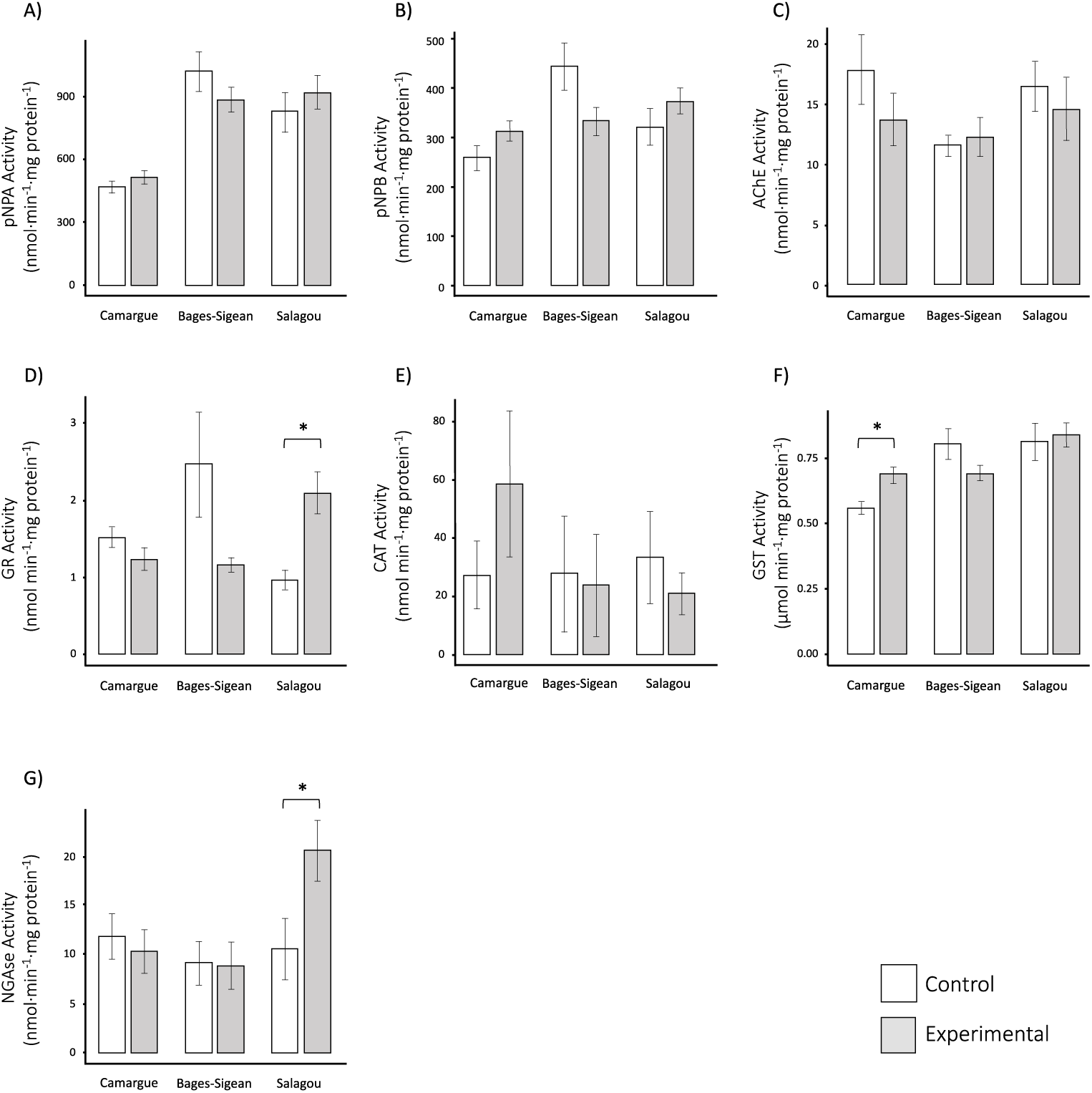
Activity of carboxylesterase using p-nitrophenyl acetate (p-NPA) and using p-nitrophenyl butirate (p-NPB), acetylcholinesterase (AChE), glutathione reductase (GR), catalase (CAT), glutathione-S-transferase (GST), and NAGase (NAG) in hepatopancreas for each of the 3 populations considered. Error bars represent standard deviation. The * represents statistically significant values at p ≤ 0.05.

**Figure 6.**
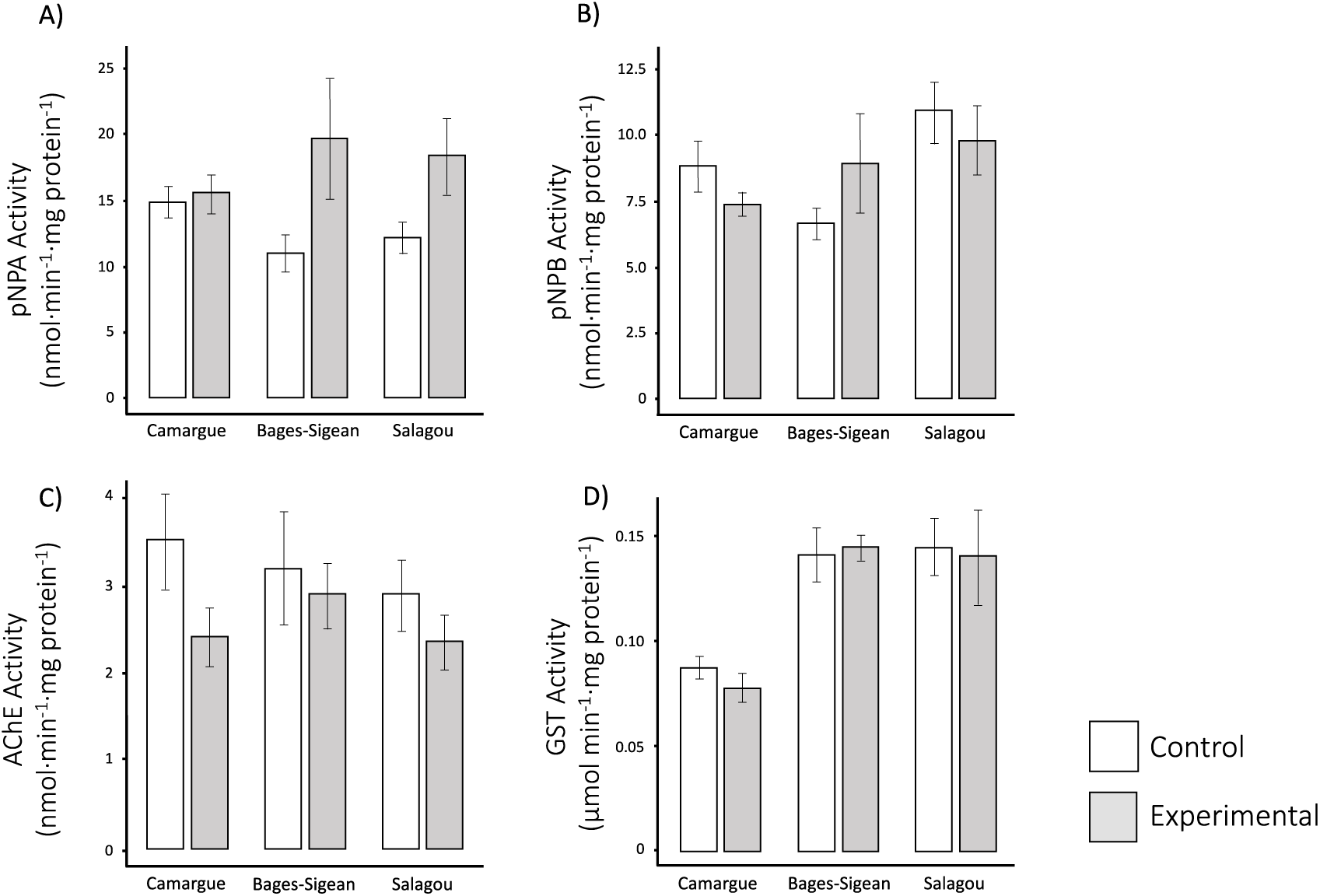
Activity of carboxylesterase (CE) using p-nitrophenyl acetate (p-NPA) and using p-nitrophenyl butyrate (p-NPB) as substrates, acetylcholinesterase (AChE) and glutathione-S-transferase (GST) in gills for each of the 3 populations considered. Error bars represent standard deviation.

**Table 2:** Enzymatic results (expressed as average ± standard error of mean) for control and experimental *Procambarus clarkii* coming from three populations in the south of France. Catalase (CAT, nmol min^−1^·mg protein^−1^), glutathione reductase (GR, nmol min^−1^·mg protein^−1^), glutathione-S-transferase (GST, µmol min^−1^·mg protein^−1^), carboxylesterase (CE nmol·min^−1^·mg protein^−1^) using p-nitrophenyl acetate (pNPA) and p-nitrophenyl butirate (pNPB), acetylcholinesterase (AChE, nmol min^−1^·mg protein^−1^) and N-Acetyl-β-d-glucosaminidase (NGAse, nmol·min^−1^·mg protein^−1^). * p ≤ 0.05, ** p ≤ 0.01, n.a.: not available.

|  |  | **Hepatopancreas** |  |  | **Gills** |  |  |
| --- | --- | --- | --- | --- | --- | --- | --- |
|  |  | Camargue | Bages-Sigean | Salagou | Camargue | Bages-Sigean | Salagou |
| **pNPA-CE** | Control | 466.2±29.7 | 1019.8±88.9 | 823.2±94.8 | 14.92±1.28 | 10.96±1.44 | 12.19±1.25 |
| | Polluted | 514.8±34.9 | 885.2±56.4 | 920.6±79.9 | 15.52±1.53 | 18.57±4.18 | 18.32±2.93 |
| **pNPB-CE** | Control | 259.9±24.0 | 443.9±45.6 | 323.9±36.7 | 8.80±0.97 | 6.64±0.62 | 10.88±1.18 |
| | Polluted | 313.5±20.0 | 333.8±26.8 | 375.5±26.9 | 7.36±0.42 | 8.74±1.66 | 9.80±1.32 |
| **AChE** | Control | 18.0±2.9 | 11.6±0.9 | 16.6±2.1 | 3.52±0.55 | 3.14±0.56 | 2.90±0.41 |
| | Polluted | 13.8±2.1 | 12.4±1.5 | 14.7±2.6 | 2.43±0.34 | 2.70±0.39 | 2.35±0.31 |
| **GR** | Control | 1.53±0.13 | 2.5±0.64 | 0.97±0.12 | n.a. | n.a. | n.a. |
| | Polluted | 1.24±0.14 | 1.16±0.09 | 2.1±0.27** | n.a. | n.a. | n.a. |
| **CAT** | Control | 27.7±11.7 | 28.0±18.6 | 33.7±15.8 | n.a. | n.a. | n.a. |
| | Polluted | 58.9±25.0 | 24.0±16.4 | 20.9±7.2 | n.a. | n.a. | n.a. |
| **GST** | Control | 0.56±0.02 | 0.80±0.05 | 0.81±0.07 | 0.09±0.00 | 0.14±0.01 | 0.15±0.01 |
| | Polluted | 0.68±0.03* | 0.69±0.03 | 0.84±0.04 | 0.08±0.00 | 0.15±0.00 | 0.14±0.02 |
| **NGase** | Control | 11.8±2.3 | 9.08±2.1 | 10.6±3.1 | n.a. | n.a. | n.a. |
| | Polluted | 10.3±2.2 | 8.8±2.2 | 20.5±3.0* | n.a. | n.a. | n.a. |

Regarding antioxidant activities (Fig. 5D, E, F), only the antioxidant enzyme GR responded to pesticide mixture exposure with Salagou animals (originally collected from the pristine freshwater site) showing a 2-fold higher GR activity than the other two more polluted populations (Fig. 5D). By contrast, detoxification phase II GST (Fig. 5F), only showed significant changes in those animals collected from Camargue (i.e. pesticide polluted site): when exposed to the mixture under laboratory conditions, Camargue animals responded by increasing their GST activity by 20%.

NAGase activity (Fig. 5G) was similar in all control animals regardless of their origin with an average of 11.07±7.3 nmol·min^−1^·mg protein^−1^. However, when experimentally exposed to pesticides for 96h, Salagou animals showed a 2-fold increase in NAGase activity, while crayfish from Camargue and Bages-Sigean decreased their activities by 17 and 20%, respectively.

Overall, crayfish from Bages-Sigean showed a decrease in pNPA-CE and pNPB-CE and GST activities in hepatopancreas when exposed to the pesticide mixture for 96h while specimens from Camargue and Salagou showed an increase in these detoxification activities. Crayfish from Bages-Sigean experienced an increase in AChE activity while the other two populations showed a decrease in activity under laboratory exposure to the pesticide mixture.

## 4. Discussion

Aquatic organisms are exposed to harmful chemicals and selection pressures associated with these exposures have led to the evolution of tolerance levels, a phenomenon that has been well-addressed in insects and their resistance to pesticides (reviewed by Hawkins et al. 2018). However, the effects of anthropogenic stressors, such as pesticide inputs, on the plasticity of invasive aquatic species has not yet been addressed. Under laboratory conditions, we here addressed this subject using a transcriptomic and metabolic approach to determine the differential response of three populations of the invasive crayfish *P. clarkii* (differing in their environmental quality and historical background of chemical pollution) to a mix of two largely used pesticides (i.e. azoxystrobin and oxadiazon).

### 4.1 Differential gene expression analysis revealed a population-specific response

The most significant result derived from the differential gene expression analysis is that the response to the pesticide mix was highly population-dependent. In this study, we hypothesized that when chronically exposed to a certain extent of environmental pollutant loads, crayfish would exhibit particular responses upon facing a new chemical challenge. Consequently, we anticipated that the Camargue and Bages-Sigean crayfish populations would show a distinct response to those from the Salagou, likely to be more vulnerable as they were less exposed to anthropogenic pollutants. Our results corroborated this hypothesis; however, they also revealed a very distinct response pattern between the Camargue and Bages-Sigean crayfish populations.

The response of the Camargue crayfish (seasonally exposed to pesticide inputs) was by far, the most pronounced, mostly characterized by a down-regulation of genes in gills, and both up-and down-regulation in the hepatopancreas, as a consequence of a new pesticide challenge. A high number of down-regulated stress genes has been observed in crustaceans exposed to cadmium (Liu X et al. 2021) and low temperatures (Yang et al. 2022). Former studies have contrasted gene expression levels between natural populations of killifish from polluted and unpolluted sites (e.g. Fisher and Oleksiak, 2007) or, in response to heavy metal cadmium (Cd^2+^) in a laboratory scenario (e.g. Liu X et al. 2021). However, to the best of our knowledge, this is the first study to address differential gene expression in populations from differentially polluted natural sites upon new pesticide load under laboratory conditions.

### 4.2 Tissue-specific response

Crayfish from Salagou (reference site) exhibited a higher DEG response in gills than in hepatopancreas while those from Camargue showed similar levels of DEGs in both tissues. Also, contrary to Camargue crayfish, the response of Salagou gills was mainly up-regulation. Likewise, the number of modulated DEGs in the freshwater prawn *Macrobrachium rosenbergii* upon Cd exposure was significantly higher in the gills than in the hepatic tissue (Liu X et al. 2021). Moreover, Liu X et al. also reported that with over exposure time the number of DEGs in gills decreased while in the hepatopancreas increased. These results suggest that the gills act as the initial site and transient storage organ during short-term exposure, with the contaminant gradually being transferred from the gills to the hepatopancreas. Following this premise, it can be anticipated that crayfish from Camargue and Salagou populations may metabolize pesticide inputs at different rates, leading to delayed hepatopancreas responses after 96 hours of exposure in the most traditionally exposed Camargue population. This is partially confirmed by the fast and significant response in NGAase and GR activities in crayfish from the reference site upon pesticide exposure. However, in order to fully validate this hypothesis, it would be necessary to follow the responses in a larger time frame in both crayfish populations. Another differential trait between these two contrasted populations is that Camargue crayfish showed in gills more down-regulated genes than up-regulated, contrary to Salagou in which most of the DGEs were up-regulated. This could also be related to the rate at which gills, the first organ exposed to these pollutants, responded.

### 4.3 Expression of defense/stress-related genes

Although the Camargue crayfish population showed the highest differential gene expression in hepatopancreas and gills, in terms of enzymatic responses, only the detoxification GST activity was statistically enhanced in hepatopancreas. These results were confirmed with the annotation of the DEGs. The particular antioxidant and detoxification gene expression and enzymatic response did not increase as expected, however, we demonstrated that other important defense mechanisms at the gene expression level such as stress response, DNA repair, immune response, and translation and transcription processes, were altered. The Camargue crayfish population differentially responded with a large number of upregulated genes in the hepatopancreas when exposed to the pesticide mixture under laboratory conditions, particularly those involved in stress response, DNA repair, immune response, and translation and transcription, while in the gills, these processes are mainly downregulated. These results also suggest that although both tissues of Camargue crayfish are highly regulated, the midgut gland upregulates genes involved in stress response, DNA repair, and immune response, while the gills downregulate them. In contrast, Salagou individuals, although also exhibiting a high response in these metabolic processes, they took place primarily in the gills, and they are mostly upregulated.

Interestingly, out of the three crayfish populations considered, only those from the pristine site (i.e. Salagou) did not show an induction of DNA repair-related genes in response to pesticide exposure in the laboratory. Normally, the activation of DNA-damage-inducible genes might be expected to confer protection preventing genotoxicity (Papathanasiou and Fornace, 1991), therefore we assume that this is their role in the case of the Camargue and Bages-Sigean crayfish and it could be seen as an adaptation to the regularly periodical exposure to pesticides that they face in their natural environment. We attribute this response of DNA damage-inducible related genes to the presence of oxadiazon in the pesticide cocktail, given it demonstrated genotoxic potential even at low concentrations (Zanjani et al. 2017). In aquatic species, regularly exposed to harmful levels of chemicals, some of the potential adaptive mechanisms of defense include enhancement of anti-oxidant responses, and increased capacity for DNA and tissue repair (Hamilton et al. 2017).

### 4.4 Expression of chitinase-related genes with a possible role in the immune response

Our results on the regulation of the chitinase or chitin related genes are also worth highlighting. These were highly downregulated in the gills of Camargue crayfish (pesticide polluted) but highly upregulated in the gills of Salagou crayfish (reference site). This enzyme not only degrades chitin during growth, development and the molting processes in arthropods, but it also plays a key role in the immune responses and regulation (e.g. Niu et al. 2018; Liu M et al. 2021). In other crustaceans like the tiger shrimp (*Penaeus monodon*) the expression of chitin genes has been detected in several tissues although the highest levels have been found in gills and hepatopancreas. It has been suggested that some chitinase-related genes may be involved in the innate immune responses in *P. clarkii* by modulating the toll pathway (Liu M et al. 2021). Furthermore, chitinase in crustacean species has been observed to strongly respond to cadmium stress (Yang et al. 2024), and gene expression levels significantly increased under ammonia-N stress (Zhou et al. 2018). As mentioned previously, we found, here, chitinase genes differentially expressed in the gills of Camargue and Salagou crayfish. In Camargue crayfish, 2 chitinase genes are down-regulated in the gills but, in Salagou crayfish they were up-regulated. It is particularly interesting that the latter were 300 and 87 times more expressed in the pesticide exposed animals than in controls, which makes it a very sensitive response. In fact, this gene upregulation is also confirmed at the enzymatic level by a 2-fold increase in NGAase activity, an enzyme involved in chitin degradation during the molting process (Rollin et al., 2021). Of the experimental crayfish, none of them were in their molting period during or after the experimental exposure took place, leading us to believe that this response is not due to the natural molting process. Hence, we attribute the chitinase response in gills to an immune response triggered by the pesticide exposure. The up regulation of this protein is particularly significant in Salagou individuals, which also exhibited a significant up-regulation of pseudo hemocyanin-like proteins, but in the hepatopancreas. Hemocyanin-like proteins are crucial immune proteins in arthropods (Decker and Jaenicke, 2004; Yan et al. 2011), and their expression has been shown to increase significantly following cadmium exposure (Liu X et al. 2021). These findings further support the hypothesis that individuals from Salagou (reference site) initiate an immune response in the gills and hepatopancreas, even if this entails a distinct set of proteins compared to those shown in crayfish from the pesticide-exposed site of Camargue.

### 4.5 Enzymatic responses: effects on antioxidant defenses, detoxification processes and health biomarkers upon exposure to a pesticide mixture

It is well recognized that the presence of xenobiotics, including pesticides, induces the production of reactive oxygen and nitrogen species (RONS), compromising antioxidant defenses and, eventually, leading to oxidative stress (e.g. Sule et al. 2022). In this study, as a consequence of a 96h exposure to the pesticide mixture, only crayfish from the Camargue population displayed an increase in GST activity in hepatopancreas. In a former study by Uçkun et al. (2021), exposure to one of the two pesticides (azoxystrobin) caused enhanced GST activity in another crayfish species. An interesting work by Kovačević et al. (2023) shows the temporal dynamics of exposure to azoxystrobin in a terrestrial invertebrate. Although the study by Kovačević et al. considered pollutant concentrations per kilogram of soil, their lowest concentrations could be comparable to the concentrations used here per liter of water (despite different bioavailability). They observed that CAT and GST activities increased during the first three days of exposure, but then decreased from day five onwards. However, most reports agree that exposure to herbicides causes a decrease in antioxidant defenses leading to oxidative stress.

CEs are potential biomarkers of pesticide exposure, as they are an important family of enzymes involved in the metabolism of xenobiotic and endogenous compounds for a wide variety of organisms, including crustaceans (Wheelock et al. 2008; Nos et al. 2021). In this study, we did not find a significant change in CE activities in the hepatopancreas or gills with either of the two substrates used. However, since the hydrolysis rates with pNPA substrate exceeds that of pNPB by approx. 50% in almost all cases, it is likely that pNPA is more adequate substrate for CE measurements in this crayfish species.

In this study, the polluted populations of Bages-Sigean and Camargue, responded to pesticide exposure in the laboratory by decreasing their NGAse activity. Existing literature that indicates that pollutants such as heavy metals (i.e. Cadmium or Zinc) (e.g. Mesquita et al., 2015; Rollin et al. 2023), herbicides (e.g. Glyphosate) or drugs (e.g. pentoxifylline) cause a decrease in NAGase activity in crustaceans (Rollin et al. 2023). Also, it has been also observed in the European green crab *Carcinus maenas* that NAGase activity decreases as a response to heavy metals (Cd in particular) and this response is dependent on the quality of their natural environment. In this sense, *C. maenas* from a relatively pristine site, shows NAGase activity inhibition when exposed to Cd while crabs from a moderately polluted site do not. Our results do not align with those with *C. maenas*, as Salagou crayfish (more pristine area) experienced a 2-fold increase in NAGase activity when exposed to the pesticide input. Likewise, Rollin et al. (2023) observed a 35% increase as a response to the fungicide dithiocarbamate in the marine prawn *Palaeomon serratus.* All in all, even if *increases* in NAGase activity as a response to pollutants are not unheard of, decreases in activity rates have been most frequently documented.

In the present study, we did not observe a significant inhibition of the AChE activity in either gills or hepatopancreas. Contrasting with our results, another crayfish species, *Astacus leptodactylus,* experienced an increase in AChE activity upon exposure to azoxystrobin (Uçkun et al. 2021). Although the role of AChE in innerved tissues is mostly neural transmission, in others it can have an immune role as acetylcholine is a signaling molecule initiating this process (Giordani et al. 2023).

## 5. Conclusions

Our results showed population-specific responses to pesticide stress in populations with different chemical backgrounds. In populations accustomed to dealing with pesticides, DNA repair, immune response, and translation and transcription mechanisms seem to have a crucial role in coping with pesticide stress. This difference also contrasted between the hepatopancreas and gills of crayfish from the studied populations. Our results highlight the importance of considering physiological plasticity and tolerance to pollutants as being population-specific, especially for risk assessments of invasive species. These population-specific responses suggest a rapid evolutionary influence of anthropogenic stressors that is well worth further investigation.

## Acknowledgments

The authors acknowledge the support of “Parc naturel regional de la Narbonnaise en Méditerranée” and “Tour du Valat” for facilitate the sampling of the individuals. We would also like to thank Sergi Omedes for his technical assistance during enzymatic analysis.

## Author contribution

<u>Martinez-Alarcon:</u> Conceptualization, Funding acquisition, Investigation, Methodology, Data curation, Formal analysis, Project administration, Supervision, Writing of original draft.

<u>Lignot:</u> Conceptualization, Investigation, Review and editing

<u>Reisser:</u> Methodology, Formal analysis, Review and editing.

<u>Solé:</u> Investigation, Methodology, Data curation, Formal analysis, Review and editing.

<u>Rivera-Ingraham:</u> Investigation, Methodology, Data curation, Formal analysis, Review and editing.

## Data availability

The datasets presented in this study can be found in online repositories. And will be released when the manuscript is accepted

## Ethics statements

No permits were required to conduct this study. However, it was conducted in accordance with the local legislation and institutional requirements. All animals were captured, manipulated and euthanized following the International Guiding Principles for Biomedical Research Involving Animals, issued by the Council for the International Organizations of Medical Sciences.

